# Predicting growth conditions from internal metabolic fluxes in an *in-Silico* model of *E. Coli*

**DOI:** 10.1101/002287

**Authors:** Viswanadham Sridhara, Austin G. Meyer, Piyush Rai, Jeffrey E. Barrick, Pradeep Ravikumar, Daniel Segrè, Claus O. Wilke

## Abstract

A widely studied problem in systems biology is to predict bacterial phenotype from growth conditions, using mechanistic models such as flux balance analysis (FBA). However, the inverse prediction of growth conditions from phenotype is rarely considered. Here we develop a computational framework to carry out this inverse prediction on a computational model of bacterial metabolism. We use FBA to calculate bacterial phenotypes from growth conditions in *E. coli*, and then we assess how accurately we can predict the original growth conditions from the phenotypes. Prediction is carried out via regularized multinomial regression. Our analysis provides several important physiological and statistical insights. First, we show that by analyzing metabolic end products we can consistently predict growth conditions. Second, prediction is reliable even in the presence of small amounts of impurities. Third, flux through a relatively small number of reactions per growth source (~10) is sufficient for accurate prediction. Fourth, combining the predictions from two separate models, one trained only on carbon sources and one only on nitrogen sources, performs better than models trained to perform joint prediction. Finally, that separate predictions perform better than a more sophisticated joint prediction scheme suggests that carbon and nitrogen utilization pathways, despite jointly affecting cellular growth, may be fairly decoupled in terms of their dependence on specific assortments of molecular precursors.

## 1 Introduction

Research into metabolism and physiology generally tries to uncover how an organism’s internal state is determined by the environment to which the organism is exposed. For example, one might ask which genes are up or downregulated as microbes are grown on different nutrient sources [1–3]. Similarly, one might ask how changes in nutrient availability and gene expression alter flux of metabolites through the organism’s metabolic network [4–7]. One can also pose the inverse question, however: If we know an organism’s physiological state, can we infer the environment in which the organism is living or was grown? In some cases *in vivo*, for example, it may be easier to measure the organism’s physiological state (as defined, for example, by gene expression levels) than correctly identifying the nutrients that the bacteria feed on. In such cases, one might want to predict the bacterial growth conditions from the measured bacterial physiology.

Do we expect physiology, and in particular the internal metabolic fluxes, to be predictive of the current environment? On the one hand, one could envision a scenario where an organism can only assume a small number of distinct metabolic states, and many diverse environments elicit the same physiological response. In other words, the mapping from environment to metabolism is many-to-one. Under this scenario, metabolism would not be particularly predictive of environment. On the other hand, each environment might elicit an entirely different metabolic response, i.e., the mapping from environment to metabolism is one-to-one. Under this scenario, organismal physiology can be considered an accurate reflection of the specific environment the organism resides in, and the environment can be predicted accurately from the metabolic state. In reality, we can expect the mapping between environment and metabolism to fall somewhere between these two extremes. While there are probably many different metabolic states an organism can assume, there will also be distinct environments that create similar metabolic responses.

Metabolic modeling approaches generally ask the forward question, i.e., how can we calculate the metabolic state as a function of the environment. For example, flux balance approaches calculate the metabolic fluxes in an organism as a function of input fluxes and the organism’s metabolic network [8–11]. More sophisticated approaches might use kinetic or dynamic models [12,13]. All these modeling approaches are mechanistic approaches that mimic the chain of causality in metabolism: environment and genetic architecture are given, and metabolic state follows from the laws of physics and biochemistry. To ask the inverse question, which metabolism corresponds to which environment, we have to go against the chain of causality. Therefore, a mechanistic model is not the most appropriate to ask this question. Instead, it makes more sense to use a statistical approach to search for associations between metabolic states and environmental conditions.

Here, we asked whether the internal metabolic fluxes in an *in-silico* model of *E. coli* can predict the organism’s simulated growth environment. We first calculated metabolic fluxes via FBA for a variety of environmental growth conditions. We then developed statistical models to predict the particular growth conditions for a particular solution of the flux balance equation, using internal metabolic fluxes as input. We found that prediction is possible with surprisingly low error rates, even if a moderate amount of impurity is present in the simulated growth media. We further found that carbon and nitrogen metabolisms seem to be largely decoupled (prediction accuracy of a given carbon source does not strongly depend on the presence of particular nitrogen sources and vice versa) and that most carbon and nitrogen sources can be identified reliably from a small number of predictive fluxes.

## 2 Results

### 2.1 Predicting growth conditions from simulated flux in E. Coli

Our overarching question is to what extent the internal metabolic fluxes in an *in-silico* model of *E. coli* encode the growth substrates. Are there distinct metabolic states that reflect specific growth conditions? We addressed this question using the following strategy (Figure 1): (i) simulate metabolic fluxes under a variety of different growth conditions (primarily distinct carbon and nitrogen sources); (ii) develop regression models that relate growth conditions to the calculated metabolic fluxes; (iii) evaluate how accurately the regression model can predict growth conditions from fluxes, and identify the most predictive fluxes.

**Figure 1.**
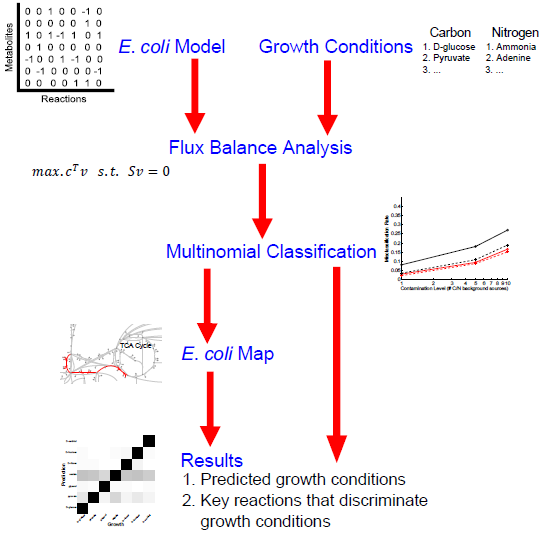
Flowchart outlining our analysis approach. We use the *E. coli* FBA model iAF1260 to generate metabolic fluxes under a variety of conditions. We then use multinomial classificaton via regularized regression to develop a model that can predict the chosen growth conditions from the metabolic fluxes. We find that prediction works generally well, even when there are chemical impurities in the growth environment. We further find that most growth conditions can be identified based on a small number of key reactions.

A biochemical network can be treated as a system that takes up the nutrients from the environment and converts them into useful metabolic precursors such as amino acids, nucleotides, and lipids. These environmental nutrients are brought into the cell via *exchange reactions*, which simply take up a molecule of a specific metabolite and place it into cell. Any metabolic flux model contains a substantial number of such exchange reactions. Within a cell, the metabolites are transferred via *transport reactions*. Clearly, predicting environmental growth conditions from fluxes through these exchange or transport reactions would be trivial, and it would not be a reflection of what information the internal metabolic state of the cell holds about the external environment. To address this issue, we discarded all transport and exchange reactions in our regression analysis. In our model (the iAF1260 metabolic model of the *E. coli* K-12 MG1655 strain [14]), this amounted to 939 reactions among a total of 2382 distinct reactions. We also discarded the biomass composition reaction, and thus were left with a total of 1442 reactions for regression analyses.

Further, to make the task of predicting growth conditions from fluxes more difficult and more realistic, we introduced background impurities in all simulated environments. Each environment consisted of a set of primary metabolites (usually one carbon and one nitrogen source) plus a small quantity of randomly chosen other metabolites. We varied the number of metabolites serving as impurities to evaluate how sensitive the regression model was to the amount of chemical noise present in the environment. Impurities were selected at random from a set of 174 carbon and 78 nitrogen sources used previously with the *E. coli* model [9]. A different set of random impurities was chosen for each individual FBA calculation. We set the maximum uptake rate of individual impurities to 1/100th of the maximum uptake rate for the main growth sources, and we tuned the amount of chemical noise in our simulations by changing the number of impurities. Throughout this work, we refer to background chemical noise of *n* carbon sources and *n* nitrogen sources as *n* C/N impurities. (We generally chose an equal number of nitrogen and carbon sources as impurities, except in two analyses where we used either only carbon or only nitrogen sources as impurities.)

We first wanted to test how well prediction might perform in a best-case scenario. To this end, we selected seven carbon and seven nitrogen sources (Table 1) that generated substantially distinct flux patterns in the absence of impurities. We assessed the similarity of flux patterns by k-means clustering of fluxes obtained for all 174 carbon and 78 nitrogen sources (see online data repository). We simulated fluxes for environments containing all pairwise combinations of the seven carbon and seven nitrogen sources, plus impurities. For each amount of impurity and training data-set size, we generated 100 replicates of each pairwise combination, for a total of 4900 sets of flux values (see Table 2 for an overview of all simulations performed). We discarded solutions that we considered to be non-viable (see Methods). We subdivided the remaining sets of flux values into two groups, a training data set and a test data set. We then fit a regularized regression model to the training data set and subsequently evaluated how well the model could predict growth conditions from fluxes on the test data set.

**Table 1.** Substrates used as primary carbon and nitrogen sources.

| **Carbon sources** | **Nitrogen sources** |
| --- | --- |
| D-glucose | Ammonia |
| Pyruvate | Adenine |
| Glycerol | Cytidine |
| Acetate | Putrescine |
| D-ribose | L-glycine |
| D-fructose | L-alanine |
| D-sorbitol | L-glutamine |

**Table 2.**
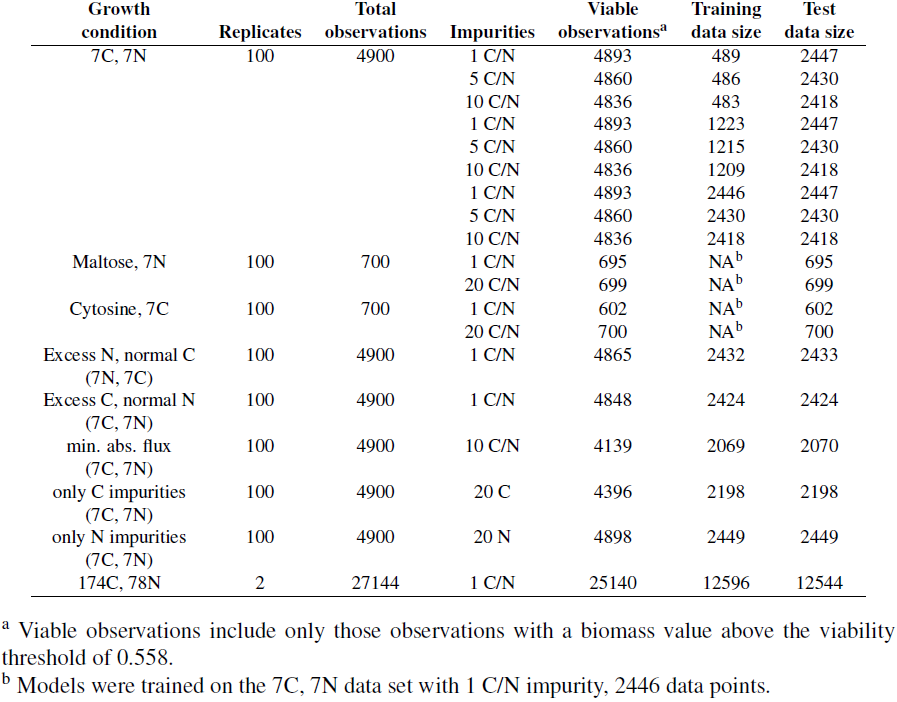
Summary of all simulations performed. Each row details the growth conditions used for flux balance analysis (FBA) and the sizes of training and test data sets for inverse prediction of growth conditions from simulated phenotypes.

We considered two alternative approaches to prediction, joint prediction and separate prediction. Under joint prediction, we considered all 49 pairwise combinations of the seven carbon and seven nitrogen sources as distinct outcomes, and we trained a single model to predict one of those 49 possibilities. Under separate prediction, we trained two separate models, one for the seven carbon sources and one for the seven nitrogen sources. Overall, both prediction approaches worked quite well. Even at relatively high numbers of impurities, we could correctly identify the main carbon and nitrogen sources in over 80% of the cases (Figure 2). And for very few impurities, i.e. 1 C/N, the misclassification rate fell below 5%. Note that by random chance, we would expect a correct prediction only one time out of 49, i.e., by random chance the misclassification rate would be 98%.

**Figure 2.**
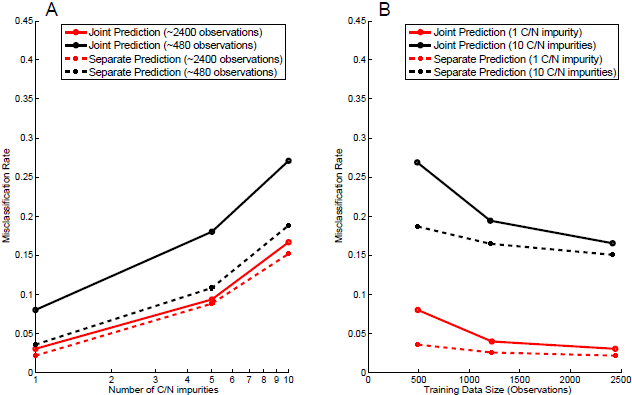
Misclassification rate versus number of impurities and amount of training data. For joint prediction, each data point corresponds to training/testing a new regression model. Similarly for separate prediction, each data point corresponds to training/testing two separate new regression models. (A) The misclassification rate increases as the number of impurities increases. (B) The misclassification rate decreases as the size of the available training data increases. In all cases, separate prediction out-performs joint prediction.

To understand where the misclassifications are coming from, we plotted heatmaps that show the actual growth sources and the predicted sources at two different numbers of impurities (1 C/N and 10 C/N). At 10 C/N, a number of carbon sources are predicted as either acetate or pyruvate (Figure 3). A closer look at the key reactions unique to these sources revealed that the reactions either are near the site of entry into the TCA cycle or within the TCA cycle. The role of TCA cycle is to generate energy, precursors for amino acids, and cofactors such as NADH. This means that under any environmental conditions, there may need to be some flux in the reactions that enter the TCA cycle. So, as the number of distinct impurities increases, the flux resulting from these impurities is seen through these common reactions and hence observations get mispredicted as acetate or pyruvate.

**Figure 3.**
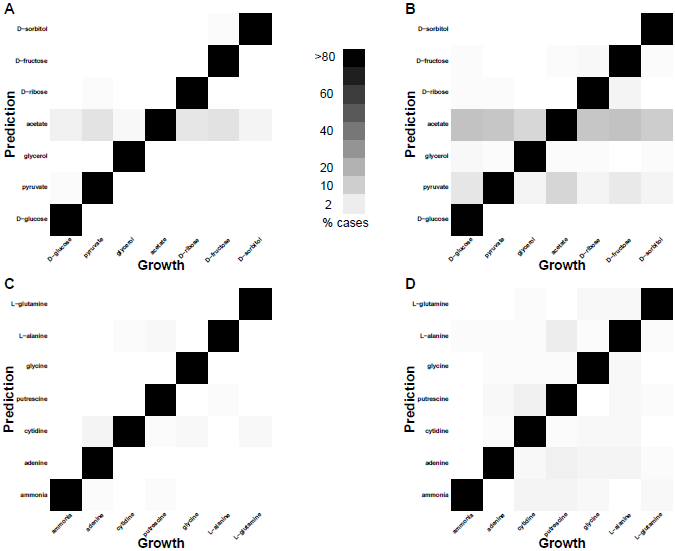
Probability of misclassification of C and N sources. For each heat map, the actual C or N source is plotted along the *x*-axis and the predicted one is plotted along the *y*-axis. The gray level of squares indicates the fraction of times a given C or N source was predicted, with white corresponding to 0% and black corresponding to 100%. (A, B) C sources predicted from models with 1 and 10 C/N impurities, respectively. (C, D) N sources predicted from models with 1 and 10 C/N impurities, respectively. In most cases, prediction is accurate (near-black squares along the diagonal). Prediction accuracy declines with the number of C/N impurities, as expected. For C sources, most misclassifications lead to a prediction of acetate, or, to a lesser degree, pyruvate. A similar pattern does not exist for N sources.

In a direct comparison, however, the separate prediction models always outperfomed the joint prediction models (Figure 2). The performance gap was virtually independent of the number of impurities, but it did depend more strongly on the size of the training data set. In particular for smaller training-set sizes, independent prediction performed much better. We assume that the advantage at small sizes of training data sets arose because the independent prediction had effectively seven times more data to train than the joint prediction. For example, if the training data set was so small that it contained only one observation for each of the 49 joint conditions, it couldn’t be used at all to train the joint model. However, two independent models (either carbon sources only or nitrogen sources only), there would be seven observations for each of the seven carbon or nitrogen sources.

Next, we looked into understanding the role of excess resources on prediction results. Above, we used the conventional maximum uptake rate of 20 mmol gDW^−1^ hr^−1^ that is generally used for carbon and nitrogen sources in FBA studies. To determine to what extent our results depended on this choice, we artificially increased the uptake rates of the carbon source to a maximum of 1000 mmol gDW^−1^ hr^−1^ while keeping the nitrogen source at the normal rate, and vice versa. These simulations can be considered as conditions of excess carbon (when maximal carbon uptake is artificially increased) or excess nitrogen (when maximal nitrogen uptake is artificially increased). Figures S1 and S2 show the realized uptake rates for these simulations.

When predicting growth conditions from the final fluxes, we obtained similar results as before, i.e., individual prediction performed better than joint prediction. For an artificially high uptake rate for nitrogen but with a normal uptake rate for carbon sources, the misclassification rate with separate prediction was 10%, while the misclassification rate with joint prediction is 26%, at an amount of impurities of 1 C/N and with a training data size of ~2450 replicates. Separately predicting carbon resulted in 151 mispredictions compared to 109 mispredictions for nitrogen. In combination, there were 250 mispredictions using separately trained models. Joint prediction resulted in 638 mispredictions. Similarly, for artificially high uptake rates for carbon sources and normal uptake rates for nitrogen sources, the misclassification rate under separate prediction was 3.8% while the misclassification rate under joint prediction was 14%. Joint prediction resulted in 324 mispredictions. Separate prediction resulted in a combined misprediction from C and N sources of 94 mispredictions (64 C and 32 N, respectively). Clearly, prediction rates were better for separate prediction compared to joint prediction even with larger training data sizes, which was not the case for *normal* uptake rates for both sources. It seems that the more one source becomes available in excess, the more separate prediction outperforms joint prediction.

Since individual prediction seemed to work well, we next tested whether we could use this approach to predict growth conditions chosen from the comprehensive list of 174 carbon and 78 nitrogen sources. Joint prediction in this case was infeasible, since we would have had to train a model to distinguish between 174 × 78 = 13572 distinct conditions. To test independent prediction in this case, we generated simulated fluxes for all pair-wise combinations of carbon/nitrogen sources for two replicates. We used one replicate (78 carbon observations and 174 nitrogen observations respectively for each of carbon and nitrogen sources) to train the regression model and we used the second replicate to evaluate the prediction accuracy of the model. We found that the mis-classification rate for carbon sources was 86% and the misclassification rate for nitrogen sources was 37%. In combination, the two models identified the correct carbon/nitrogen combination 8.7% of the time. By random chance, we would have expected 1/13572 = 0.007%.

All the results presented so far were obtained with a simple maximization of the biomass reaction. FBA can also be carried out with different optimization functions, and the equilibrium fluxes that are found will depend on the specific optimization function chosen. To confirm whether our approach would work under different optimization schemes, we carried out additional simulations in which we maximized biomass and then subsequently minimized the absolute sum of fluxes, holding the maximal biomass value constant. Then we performed the regression analysis as described above. We carried out this analysis for the case of 7 distinct C and 7 distinct N growth substrates, 10 C/N impurities, and individual prediction of C and N sources. We found a combined misclassification rate of 23%, relative to 15% using only the biomass maximization. (See Figure S3 for prediction results for individual C and N sources.)

### 2.2 Identifying the predictive fluxes

The previous subsection has shown that a regularized regression model is capable of predicting the primary carbon and nitrogen sources used from steady-state metabolic fluxes. We next wanted to investigate how exactly the regularized regression model carries out this task. For each flux balance simulation, the resulting flux data set contains 1443 flux values, corresponding to 1443 reactions that are not transport reactions. One of these reactions is the biomass reaction, which we excluded from the regression modeling. Thus, we have 1442 predictor variables in the regression model. In this situation, a standard regression model would have to determine 1443 regression coefficients, one per reaction plus an intercept. By contrast, the regularized regression model we employed sets most regression coefficients to zero and retains only a small number of non-zero coefficients. (The exact number of non-zero coefficients is determined through the choice of a tuning parameter, which is selected by cross-validation. See Methods for details.) Thus, we can consider the fluxes with non-zero regression coefficients as *predictive fluxes.* Those are the fluxes whose state is actually used for prediction.

To gain mechanistic insight into predictive reactions, we mapped them onto the *E. coli* central metabolism (Figure 4 and Figure 5, Tables S1 and S2). Note that each of the metabolic maps is meant to highlight only the central carbon metabolism in the *E. coli* metabolic network. We found that each carbon or nitrogen source had a few reactions that were predictive of that growth source, and these reactions generally made physiological sense. For example, using acetate as the carbon source unsurprisingly isolated TCA cycle entry as a predictive reaction (Figure 4). The key reactions identified for D-glucose utilization were glucose 6-phosphate dehydrogenase (G6PDH), glucose-6-phosphate isomerase (PGI) and 6 phosphogluconolactonase (PGL), required enzymes in the glycolytic and pentose phosphate pathways [15]. Similarly, sorbitol (the singly reduced alcohol of D-glucose) and fructose each possessed predictive reactions in the relative vicinity of the glycolytic pathway (Figure 4). Mapping nitrogen sources to the central metabolism revealed a similar trend. For example, L-alanine as a growth source had predictive reactions near its site of entry into the three and four carbon metabolism of the TCA cycle (Figure 5).

**Figure 4.**
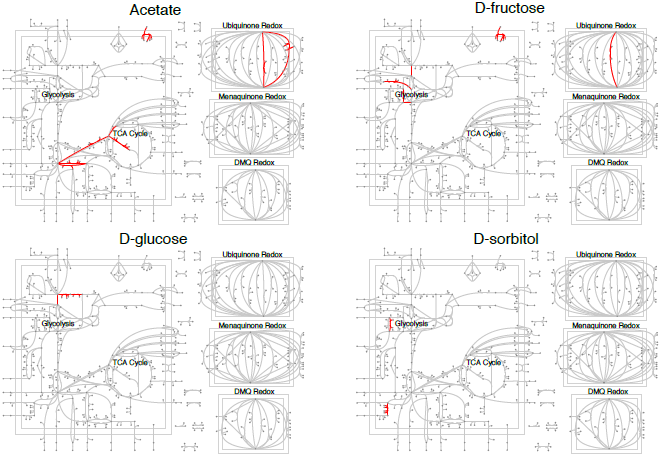
Predictive reactions (in red) for four carbon sources, mapped onto the *E. coli* central metabolism. Reactions in distinct parts of the metabolism are predictive for different carbon sources. A list of the predictive reactions can be found in Table S1.

**Figure 5.**
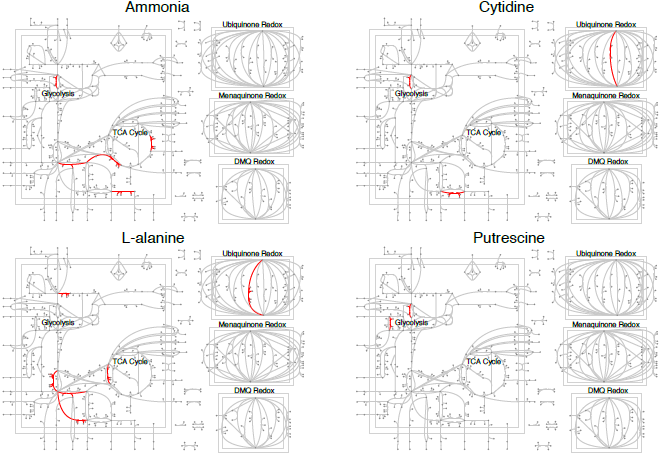
Predictive reactions (in red) for four nitrogen sources, mapped onto the *E. coli* central metabolism. Reactions in distinct parts of the metabolism are predictive for different nitrogen sources. A list of the predictive reactions can be found in Table S2

At an impurity number of 1 C/N and using the largest training data size (see Figure 2), there were 72 key reactions discriminating 7 carbon sources and 72 key reactions discriminating 7 nitrogen sources. So, on average, the regression models required 10.3 predictive reactions per carbon or nitrogen source. We found a moderate amount of overlap among the two sets of 72 reactions. 44 reactions appeared in both regression models resulting in 100 unique key reactions for 7 C and 7 N sources. Lists of the predictive reactions for each growth source are provided in Tables S1 and S2.

We also analyzed how the regression model performed when some of the key predictive reactions were removed. As mentioned above, there were 100 unique reaction IDs for individual prediction of carbon and nitrogen sources at the lowest number of impurities and with the largest training data set analyzed. We eliminated each of these 100 reactions at a time as predictors in the regression model, trained a new model separately for both the carbon and nitrogen sources, and calculated the prediction accuracy. We combined the results of individual predictions to calculate the prediction accuracy of the combination of the sources. With the exception of the reaction “glucose 6-phosphate isomerase” (PGI), the misclassification rate remained unchanged when we eliminated any of the other 99 reactions before model fitting. PGI catalyzes a reaction that produces fructose-6-P from glucose-6-P, and knock-out of the PGI gene causes diminished growth rate 16. The reaction PGI catalyzes is reversible, but the forward reaction occurs most of the time unless concentrations of fructose-6-P are very high. Thus, PGI seems to be critical to distinguish glucose from fructose as growth source. However, more generally, the specific set of reactions used for successful prediction was not unique.

Finally, we asked whether we could predict growth substrates simply on the basis of flux through the entry points of the metabolites into the metabolic network, i.e., based on first set of reactions past transport. For the 7 C and 7 N substrates we considered, there are 38 such post-transport reactions (Table 3). Thus, we built a regression model that contained only these 38 reactions as predictors for the growth substrates. At 1 C/N impurities, we obtained a combined misclassification rate of 6%, relative to 2.16% using the entire metabolic model. Similarly, at 10 C/N impurities, the combined misclassification rate was 25%, relative to 15% using the entire set of internal fluxes. This result suggests that the growth substrates are non-trivially encoded in more central parts of the metabolic network. Just knowing the flux through post-transport reactions is not as useful for predicting growth substrates as knowing the flux through the most predictive reactions, wherever they fall inside the metabolic network.

**Table 3.**
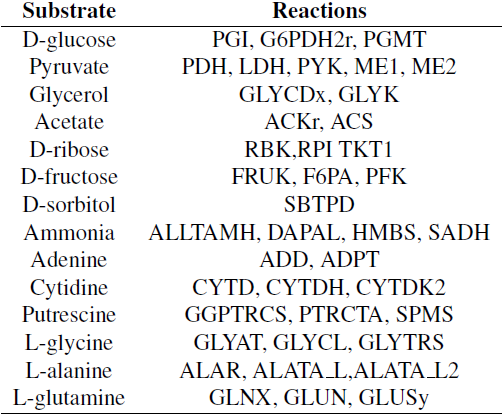
Post transport/exchange reactions that first metabolize a given substrate. Instead of using all internal reactions in the regression model, we also considered a regression model that contained only the post-transport reactions listed here.

### 2.3 Predicting novel metabolites

Next, we wanted to determine how the prediction would perform on previously unseen carbon or nitrogen sources. We first obtained simulated flux measurements using maltose as the carbon source and using either of the seven nitrogen sources used earlier. We generated simulated flux data for 100 replicates and at impurity numbers of 1 C/N and 20 C/N. This resulted in 700 observations. After eliminating replicates with very low biomass (see Methods), 695 and 700 viable flux measurements remained at 1 C/N and 20 C/N impurities, respectively. We used all these observations for testing. Note that we trained the model using the carbon/nitrogen sources in Table 1. When we tested individual prediction of either carbon or nitrogen sources, we found that maltose was classified as glucose over 85% of the time. Since maltose is a disaccharide formed from two units of glucose, this prediction is reasonable. At the same time, the seven nitrogen sources were predicted correctly over 95% of the time. However, when we tried to predict using the joint model, we found that using an unknown carbon source had a substantial effect on the model’s ability to predict nitrogen sources. 33% of the growth conditions (231 cases) were predicted to be sorbitol/putrescine. Sorbitol is a reasonable choice considering the model had never seen maltose. (Sorbitol is the singly reduced alcohol of glucose.) However, of the 231 cases, only 98 had actually been grown on putrescine, so the joint model’s prediction of the nitrogen source was misled by the unknown carbon source.

For 20 C/N impurities, there were 699 viable flux measurements. At this amount of chemical noise, maltose was predicted as glucose 68% of the time, while the correct nitrogen source was predicted 81% of the time. For both low and high numbers of impurities, individual predictions seem to outperform joint prediction. Further, separate prediction is more likely to correctly predict all the known growth sources while predicting the unknown ones to their nearest known compound.

Next, we did simulations to test how an unseen nitrogen source gets predicted with the above models. For this, we used cytosine as a nitrogen source and either of the 7 carbon sources used earlier. Note that cytosine is one of the 4 bases founds in DNA and RNA. We used 2 numbers of impurities, 1 C/N and 20 C/N, and we generated 100 replicates for each case for testing. At 1 C/N, there were 602 viable flux measurements, i.e., for these measurements biomass was greater than the threshold used in this study. Interestingly, all 98 non-viable flux measurements were for Cytosine + Acetate sources. For the viable flux measurements, only 5 carbon sources were wrongly predicted (~ 0.01% misclassification). Interestingly, in all cases, the nitrogen source cytosine was predicted as ammonia. This result may be due to a reaction that directly liberates the exocyclic amine of cytosine as ammonia.

At 20 C/N impurities, all the 700 flux measurements were viable (biomass greater than threshold). In this case, 27 carbon sources were incorrectly predicted (~ 0.04% misclassification rate). The nitrogen source cytosine is predicted as ammonia in 78.8% of cases and as adenine in all other cases.

## 3 Discussion

We have developed a method for making predictions regarding bacterial growth conditions from known simulated metabolic fluxes. We generated fluxes using the complete *E. coli* metabolic network model for 7 carbon and 7 nitrogen sources. Then, we divided the data into training and test sets and employed machine learning with a generalized linear framework to train a model to predict growth conditions. We found that even in growth environments contaminated with other nutrient sources, we could make reliable predictions regarding the primary composition of the growth media. Our prediction technique worked well on pairwise combinations of carbon and nitrogen sources, and it could also extrapolate to previously unseen metabolites.

It was surprising to us that given the same number of observations in the training set, separate prediction of nutrients always performed better than joint prediction. There are two likely explanations for this result. First, making joint predictions requires discriminating between 49 different pairwise combinations. By contrast making individual predictions only requires discriminating 7 different conditions in two different sets. Thus, one possible explanation for the lack of predictive power is that we simply did not have the appropriate level of training data. Indeed adjusting the amount of training data appears to have a dramatic effect on joint prediction in particular (Figure 2). On the other hand, such an issue represents an important experimental concern. Often the size of the training set, being experimentally determined, is just as limiting as the size of the testing set. As a result, our analysis indicates employing a separate prediction strategy will generally be more useful for experimental application. Second, although the mechanism is not completely clear to us, separate prediction may gain additional power due to the physiology of the organism. For example, if the initial, metabolite-unique, steps of metabolic entry are often predictive (as they appear to be), running independent predictions would be expected to perform better per amount of data; in essence such a prediction strategy makes the assumption that pathways for the various metabolites are largely disconnected. By contrast, if one were using a single metabolite as a combined carbon and nitrogen source, we may expect an independent prediction strategy to perform relatively poorly as the independence assumption is not satisfied.

Although the background chemical noise can have a dramatic affect on model accuracy, the misclassification rate remained acceptably low even with 10 randomly picked C/N impurities. The addition of these impurities revealed one interesting and unexpected physiological hypothesis about the *E. coli* metabolism. Namely, as the environmental chemical noise increases from 1 C/N to 10 C/N, our model increasingly predicts acetate and pyruvate as the default carbon sources. Due to its centrality in terms of energy and precursor-molecule production, for any input growth source the reactions that lead to the TCA cycle or the reactions within the TCA cycle should have some reasonable amount of reaction flux. In other words, acetate and pyruvate as default nutrient sources may not be so surprising when one considers their central role in the TCA cycle—which is essentially the center of bacterial metabolism.

To verify that the observed default carbon-source misclassification was not an artifact of nutrient limitation (carbon versus nitrogen), we increased the uptake rates of carbon source artificially high while keeping nitrogen source at normal uptake and vice versa. This ensures limiting conditions for one source and non-limiting for the other. These simulations did not alter our earlier conclusion that separate prediction performs better than joint prediction.

In addition to these simulations, we carried out three further analyses. First, instead of using all the metabolic reactions in the iAF1260 model, we used only the post-transport reactions in the regression model, as the earlier analysis had suggested that the key reactions for a growth substrate seemed often to be at the substrate’s entry point into the metabolic network. However, this approach lead to poorer predictions than did the approach of initially using all metabolic reactions in the model and letting the LASSO technique select the predictive ones. This finding confirms that the growth substrates are non-trivially encoded in the internal fluxes of the metabolic network. Second, instead of using equal numbers of carbon and nitrogen impurities, we also considered using only carbon or only nitrogen impurities. When only carbon (or nitrogen) impurities were present, prediction of nitrogen (or carbon) sources had higher sensitivity than when there was a mixture of C/N impurities. Finally, we considered an alternative optimization protocol where we minimized the absolute sum of fluxes on the FBA solution obtained by maximizing biomass. Under this protocol, prediction accuracy was somewhat lowered relative to our default protocol. However, prediction remained possible at accuracies way above random guessing.

Our regression model had a relatively large feature space (1442 reactions) compared to the number of observations used to train the model (~480 to ~2450). Therefore, efficient feature reduction was crucial to obtain reliable models. We prevented over-fitting during feature selection by employing regularized regression via the LASSO [17]. This method has previously been used successfully in other biological applications, for example to predict gene regulatory networks [18], to identify SNPs in GWAS studies[19], or to classify structural images of the brain using MRI data [20–22]. Alternative statistical methods one could consider in future work include graphical LASSO [23] and Ising Markov Random Field models [24]. We chose the standard LASSO because it provides a relatively simple and particularly robust framework for feature reduction. Thus, considering the large size of our simulation model, we were able to achieve a remarkbly small number of source-predicting reactions.

Our work is conceptually related to the work by Brandes et al. [25]. They used FBA combined with measured gene-expression data to identify candidate nutrients that likely caused the observed differences in gene-expression patterns. Their approach was based on the idea that gene-expression patterns can be expressed as flux limits, and that the biomass production rate would be larger for better candidate nutrient matches to the actual nutrient on which the microbe was grown. It is well established that gene expression levels reflect environmental growth conditions in microbes [26,27]. By contrast to Brandes et al. [25], who used a metabolic-modeling approach to identifying likely nutrients, our approach to predicting the growth environment was purely statistical. Thus, we are asking whether the physiological state of an organism contains sufficient information *in principle* to make inferences about the growth environment. Our results indicate that this is indeed the case, though it remains to be tested how well our results carry over to experimental systems.

One shortcoming of our statistical approach to predicting growth conditions is that it cannot predict previously unseen nutrients, i.e., carbon or nitrogen sources that were not used in the training data set. Nevertheless, we found that our regression model made reasonable choices, such as predicting the previously unseen maltose (a disaccharide consisting of two glucose molecules) as glucose. In this context, it is comforting that separate prediction generally outperformed joint prediction, since separate prediction was much more robust to previously unseen nutrients. In particular, a previously unseen carbon source did not substantially negatively affect prediction of a previously seen nitrogen source and vice versa.

Throughout this work, we have used basic flux balance analysis to predict the bacterial phenotype. In principle, one could use more realistic models that integrate regulatory information and/or signalling-pathway information with flux balance analysis techniques [12,28]. However, our study was a pure simulation study, primarily aimed at testing to what extent a statistical approach could be used to identify growth conditions from physiological measurements. Since our results indicate that this approach is feasible, even with a relatively modest number of observations in the training data set, a more useful next step would be to try the same approach on experimental data, using measured mRNA or protein abundances as features in the regression model.

## 4 Conclusions

We have found that predicting growth conditions from simulated metabolic flux data is a computationally tractable problem. Of note, our data indicate that separately predicting carbon and nitrogen sources performs better than jointly predicting them from paired input. Although this result is to some extent influenced by the volume of training data, it very likely reflects the structure of the metabolic reactions in the *E. coli* network. In addition, our results indicate that for most input metabolites at least one predictive reaction commonly occurs near its entry point into central metabolism. Finally, we found that the number of reaction fluxes required to make accurate predictions is relatively small.

## 5 Materials and methods

### 5.1 Flux balance analysis

We carried out flux balance analysis (FBA) using the COBRA toolbox [29] for MATLAB. We used the iAF1260 model from the BiGG database [14]. In the current iAF1260 model, there are 2382 reactions involving 1668 metabolites. The biomass composition reaction is also included in the model. Except for the input growth sources (carbon and nitrogen sources), we left all parameter settings at their default for this model. The upper bounds on 2377 reactions is set to 1000 mmol gDW^−1^ hr^−1^. But for 5 reactions, i.e., ATPM, CAT, FHL, SPODM, SPODMpp, the upper bound is set to 50 mmol gDW^−1^ hr^−1^. The lower bound for the majority of reactions (nearly 1800) is set to 0 mmol gDW^−1^ hr^−1^, which means that the flux cannot flow in the direction opposite to that specified in the model (irreversibility constraint). A set non-growth associated maintenance (NGAM) of 8.39 mmol gDW^−1^ hr^−1^ is used for the ATPM reaction (ATP maintenance). The lower bounds of some exchange reactions is set to non-zero values, i.e., these reactions by default are meant to uptake ions, carbon and nitrogen sources, and so on. We used the default oxygen uptake rate (−18.5 mmol gDW^−1^ hr^−1^) in the iAF1260 model, but we changed the lower bounds of glucose and ammonia to zero, except when we used these compounds as carbon or nitrogen sources.

After setting up the constraints on transport fluxes as described in the previous paragraph, we carried out FBA using biomass as the objective function. FBA finds fluxes that are consistent with the given constraints and that maximize the objective function. The FBA was performed with the function `optimizeCbModel` in the COBRA toolbox, using default parameters. We considered all solutions with a biomass flux below a threshold of 0.558 to be inviable. The choice of this threshold value is explained in subsection “Impurities” below.

To understand if the choice of objective function affects the prediction of growth substrates, we also carried out simulations with an additional level of constraints. The additional constraint we im-posed was minimization of the absolute sum of fluxes on the solution obtained from prior biomass maximization. This analysis was carried out using the function `optimizeCbModel(model, [], ́oné)`, where `model` is the iAF1260 model with biomass reaction as optimization function.

### 5.2 Growth conditions

We initially carried out simulations on 49 growth conditions consisting of all pairwise combinations of 7 carbon and 7 nitrogen sources (Table 1). The compounds were selected from among the 174 carbon and 78 nitrogen sources previously used in Feist et. al [9], and they were chosen to yield distinct flux profiles, as assessed by *k*–means clustering of steady-state fluxes obtained using all pair-wise combinations of the 174 carbon and 78 nitrogen sources. Of the 7 carbon sources chosen, none resulted in any growth when used as a growth condition in the absence of a nitrogen source. By contrast, all nitrogen sources except ammonia yielded growth when supplied in the absence of a carbon source. Note that all of these remaining nitrogen sources do contain carbon atoms, thus additional supply of carbon was not strictly necessary for growth on these substrates.

For any given growth environment we simulated, we set the lower bound of the exchange reactions corresponding to the carbon and nitrogen sources present to −20 mmol gDW^−1^ hr^−1^.
This lower bound is commonly used in many studies [9]. We set the lower bound of all other exchange reactions for carbon and nitrogen sources to 0, except for impurities (see next subsection for details). For each of the 49 pair-wise combinations of the 7 carbon and 7 nitrogen sources, we generated 100 replicates, each with different, randomly chosen impurities. A complete overview of all simulation conditions used is given in Table 2.

For conditions with excess carbon or nitrogen, we increased the maximum uptake rate of one source while keeping the other one fixed. Thus, we changed the lower bounds (uptake rate) of carbon sources to −1000 mmol gDW^−1^ hr^−1^ while keeping the lower bounds of nitrogen sources at −20 mmol gDW^−1^ hr^−1^ and vice versa.

Finally, we also carried out simulations on all pairwise combinations of all 174 carbon and 78 nitrogen sources previously used in Feist et. al [9], again with impurities as explained below. For this analysis, we only used two replicates for each pairwise combination of carbon and nitrogen sources, one for training and one for model evaluation.

### 5.3 Impurities

To make the simulation scenario more challenging and more realistic, we incorporated different numbers of impurities (chemical noise) to the simulated growth media. For this, we used a subset of the 174 carbon and 78 nitrogen sources, previously used in Feist et. al [9]. We used different background impurity numbers, ranging from 1 C/N to 10 C/N sources. For example, if we want to set 5 C/N impurities, we randomly picked 5 carbon and 5 nitrogen sources and set their lower bounds to −0.2 mmol gDW^−1^ hr^−1^. Note that we generated the flux data for a pairwise combination of 1 carbon and 1 nitrogen source along with the background impurities as described above. We also did additional simulations where we used only C or only N impurities. The bounds on the uptake rates for these impurities was the same as before.

For all the results described above, we used a biomass threshold to filter out non-viable flux measurements. We calculated this threshold value using biomass measurements at the lowest number of impurities (1 C/N), using all pairwise combinations of the 7 carbon and 7 nitrogen sources
chosen for the first analysis, and with the largest training dataset size (~ 2450 replicates). We recorded the biomass values for all these simulations, and used as lower threshold of viability three standard deviations below the mean, which came out to 0.558.

### 5.4 Regularized regression

We predicted growth conditions using regularized multinomial logistic regression, as implemented in the GLMNET package [30] for R. Unless specified otherwise below, we used the standard settings of the GLMNET package.

After filtering for biomass, for each number of impurities, we used half of the dataset as test set. We used subsets of the remaining half as training sets (i.e, ~245, ~490, ~2450 observations). On the training sets we fitted regularized regression models via 3–fold cross validation, using the function cv.glmnet in the GLMNET package. When using this function, we set *α* = 1 to specify the least absolute shrinkage and selection operator (LASSO). We set the parameter family equal to “multinomial”. The cross-validation procedure yielded the *λ* value that had the lowest misclassification rate under cross-validation (*λ* is the tuning parameter that controls the strength of the LASSO penalty introduced) as well as the regression coefficients at that *λ*. We then used the fitted models to calculate the misclassification rate on the test set, using the function predict. We repeated this step to calculate the misclassification rates at different numbers of impurities (1 C/N, 5 C/N, 10 C/N) and different training data sizes (~ 245, ~ 490, ~ 2450 observations).

To guarantee that the LASSO model would converge, we imposed a minimum threshold of 10^-6^ on the magnitude of all flux values. Absolute flux values below the threshold were set to zero before fitting the LASSO model.

### 5.5 Raw data and analysis scripts

All raw data and analysis scripts are available online in the form of a git repository at https://github.com/clauswilke/Ecoli_FBA_input-prediction

## 6 Acknowledgments

We thank members of the Segré lab for helpful discussions on flux balance analysis. We thank Dakota Derryberry for a critical reading of the manuscript. The Bioinformatics Consulting Group and the Texas Advanced Computing Center (TACC) at UT provided high-performance computing resources.

## Supplementary Figures

**Figure S1: Scatter plot showing varying uptake amounts of C/N sources when carbon is unlimited**. We increased the upper bounds of the carbon sources and plotted the uptake amounts of carbon and nitrogen sources. The color coding reflects whether separate prediction could predict the carbon and nitrogen sources correctly for each replicate. Green: both C and N source are correctly predicted; blue: only N source is correctly predicted; purple: only C source is correctly predicted; red: neither source is correctly predicted.

**Figure S2: Scatter plot showing varying uptake amounts of C/N sources when nitrogen is unlimited**. We increased the upper bounds of the nitrogen sources and plotted the uptake amounts of carbon and nitrogen sources. The color coding reflects whether separate prediction could predict the carbon and nitrogen sources correctly for each replicate. Green: both C and N source are correctly predicted; blue: only N source is correctly predicted; purple: only C source is correctly predicted; red: neither source is correctly predicted.

**Figure S3: Probability of misclassification, for FBA with and without minimization of absolute fluxes**. For each heat map, the actual C or N source is plotted along the *x*-axis and the predicted one is plotted along the *y*-axis. The gray level of squares indicates the fraction of times a given C or N source was predicted, with white corresponding to 0% and black corresponding to 100%. (A, C) Predictions performed on fluxes whose absolute values were not minimized (default throughout this work). (B, D) Predictions performed on fluxes whose absolute values were minimized. In all cases, we added 10 C/N impurities to the growth environment.

## Supplementary Tables

**Table S1: Key reactions identified to discriminate carbon sources**. These reactions were identified from a model trained to only predict carbon sources (separate prediction). The model was trained on a data set obtained with 1 C/N impurity and with training data-set size of ~ 2450 observations.

**Table S2: Key reactions identified to discriminate nitrogen sources**. These reactions were identified from a model trained to only predict carbon sources (separate prediction). The model was trained on a data set obtained with 1 C/N impurity and with training data-set size of ~ 2450 observations.

